# Extended Recognition of the Histone H3 Tail by Histone Demethylase KDM5A

**DOI:** 10.1101/854265

**Authors:** Nektaria Petronikolou, James E. Longbotham, Danica Galonić Fujimori

## Abstract

Human lysine demethylase KDM5A is a chromatin modifying enzyme associated with transcriptional regulation due to its ability to catalyze removal of methyl groups from methylated lysine 4 of histone H3 (H3K4me3). Amplification of *KDM5A* is observed in a number of cancers, including breast cancer, prostate cancer, hepatocellular carcinoma, lung cancer and gastric cancer. In this study, we employed alanine scanning mutagenesis to investigate substrate recognition of KDM5A and identify the H3 tail residues necessary for KDM5A-catalyzed demethylation. Our data show that the H3Q5 residue is critical for substrate recognition by KDM5A. Our data also reveal that the protein-protein interactions between KDM5A and the histone H3 tail extend beyond the amino acids proximal to the substrate mark. Specifically, demethylation activity assays show that deletion or mutation of residues at positions 14-18 on the H3 tail results in an 8-fold increase in the K_M_^app^ compared to wild-type 18mer peptide, suggesting this distal epitope is important in histone engagement. Finally, we demonstrate that post-translational modifications on this distal epitope can modulate KDM5A-dependent demethylation. Our findings provide insights into H3K4-specific recognition by KDM5A as well as how chromatin context can regulate KDM5A activity and H3K4 methylation status.

---

By regulating chromatin structure and accessibility, post-translational modifications (PTMs) on histone proteins control several cellular processes such as transcription, cell differentiation and DNA damage repair.^1–2^ These modifications are dynamic as they are installed, read, and removed by specialized proteins. Histone lysine demethylases (KDMs) are enzymes that demethylate lysine residues in histone proteins. The KDM5 subfamily of the histone demethylases specifically remove methylation from lysine 4 in histone H3 (H3K4me1/2/3), a mark associated with active transcription. KDM5 family members share a conserved multi-domain architecture composed of the jumonji N and C domains (JmjN and JmjC), which together comprise the catalytic domain, a DNA binding ARID domain, a zinc-finger domain (ZF), and two-to-three plant homeodomain (PHD) reader domains (Figure 1a).

**Figure 1:**
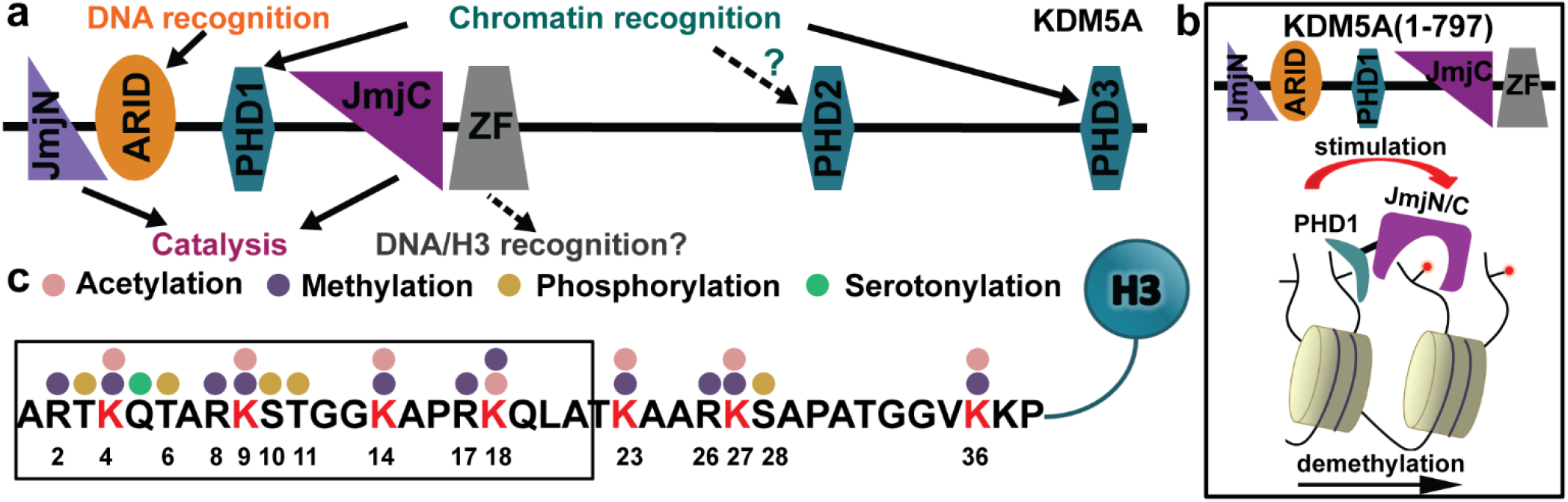
Domain architecture in KDM5A and H3 substrate sequence. **a)** KDM5A domain architecture. **b)** Proposed model for allosteric stimulation of demethylation by its PHD1 domain; the product of KDM5A-catalyzed demethylation binds to the PHD1 and stimulates catalytic activity, allowing feed-forward regulation. **c)** Sequence and select post-translational modifications of the H3 tail.

There are four paralogs of KDM5 enzymes in humans, KDM5A-D. KDM5A has three PHD reader domains (Figure 1a). The PHD1 domain binds preferentially to the product of KDM5A demethylation (H3K4me0) and allosterically enhances demethylase activity (Figure 1b).^3–4^ Function of the PHD2 is yet to be determined, while the PHD3 domain recognizes H3K4me3 marks and is postulated to recruit KDM5A to its substrate.^5^ While PHD1 is required for efficient demethylation in cells, the deletion of the PHD2 and PHD3 domains does not abrogate demethylase activity, and a KDM5A construct that lacks these domains (KDM5A_1-797_) has been utilized for the in vitro characterization of this protein (Figure 1b).^3–4, 6–7^

The presence of multiple methylated lysine residues in the H3 tail (Figure 1c) raises questions about the molecular basis for H3K4-specific demethylation by KDM5A.^8–11^ Currently, there is limited information regarding the molecular details of KDM5A substrate recognition. While KDM5 enzymes have been previously crystallized,^12–18^ none of these structures contain histone substrate. The co-crystal structure of a KDM5 plant homolog with a 10mer H3 peptide provided insights into substrate engagement by the plant enzyme,^19^ which substantially differs from human KDM5 enzymes as it lacks ARID and PHD1 domains.

In this study, we identified H3 residues necessary for substrate recognition as well as interactions between KDM5A and its peptide substrate at sites distant from the methylated H3K4 residue. To our knowledge, this is the first time that interactions between KDM5A and distal residues of H3 peptide are identified as significant for peptide binding and modulation of KDM5A activity.

## RESULTS AND DISCUSSION

### Contributions of the lysine proximal residues to K4-specific demethylation by KDM5A

To probe KDM5A-H3 substrate interactions, we investigated how alanine mutations in a H3 N-terminal 21mer (aa A1-A21) peptide substrate affect demethylase activity (Figure 2). The activity of KDM5A_1-797_ was most significantly impaired towards peptides harboring mutations in the first 9 residues, with the exception of the T3A mutant peptide (R2A, Q5A, T6A, R8A and K9A peptides) (Figure 2). Single alanine substitution of the C-terminal half of the peptide (aa 10-20) had no significant effect on KDM5A_1-797_ activity, apart from P16A and R17A mutants, which retained only half of the activity of their wild type counterpart.

**Figure 2:**
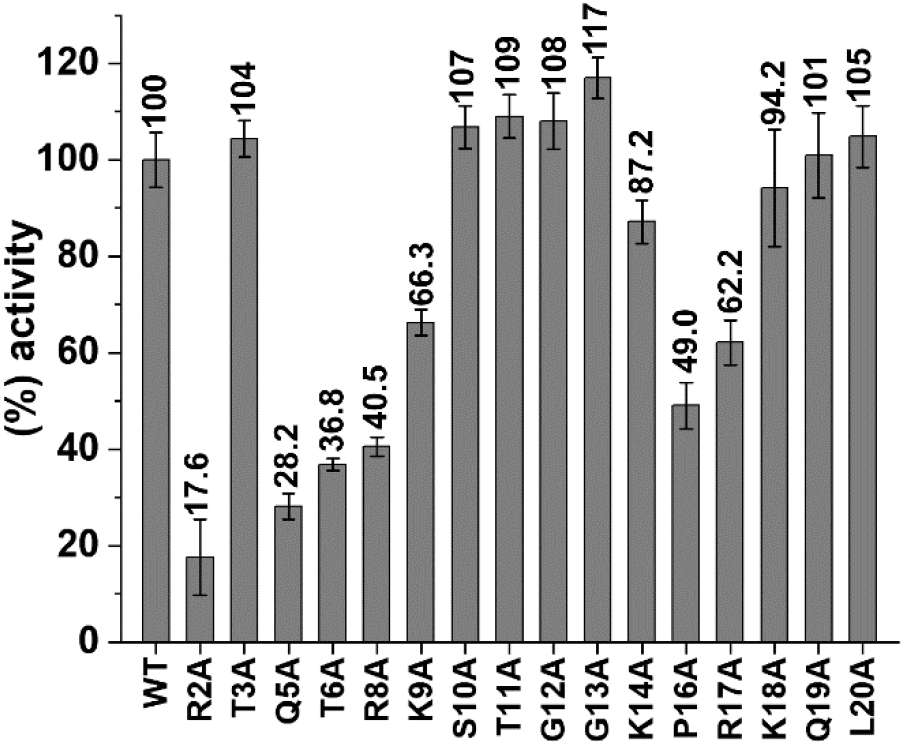
Alanine scanning mutagenesis of histone H3 tail in KDM5A-catalyzed demethylation. KDM5A_1-797_ activity for 21mer H3K4me3 WT and mutant peptides. KDM5A_1-797_ activity was normalized to the 21mer H3K4me3 WT peptide reaction (considered as 100%). Results are means ± SEM of two independent experiments.

As the first 10 residues of the H3 tail are also recognized by the PHD1 domain, and peptide binding to PHD1 stimulates demethylation activity, it is possible that the observed reduction in demethylation is due to impaired engagement of the mutant peptide by the PHD1 domain, diminishing allosteric stimulation. To test this, we pre-incubated KDM5A_1-797_ with unmodified H3K4 peptide (effector peptide) at a concentration where the PHD1 is saturated with effector peptide (20x the previously determined KD) allowing for maximal allosteric stimulation,^3–4^ and compared the activity for wild-type (WT) and mutant H3K4me3 peptide substrates (Figure 3a). In the case of R2A, T6A and R8A mutant substrates, the activity was only partially rescued in the presence of the saturating PHD1 effector peptide (Figure 3a). These findings suggest that the observed reduced demethylation of these mutant substrates is a consequence of both their decreased engagement by the catalytic domain and reduced allosteric stimulation (Figure 3a). The most prominent (~3-fold) rescue in demethylation in the presence of the effector peptide was observed with R2A mutant peptide. Indeed, our previous studies point to decreased engagement of R2A peptide by the PHD1 domain.^4^ In contrast, nearly complete rescue of K9A demethylation suggests minimal role of this residue in substrate recognition. Notably, very low KDM5A_1-797_ activity towards the Q5A substrate peptide was observed (~20%). This reduction in activity could not be rescued by the presence of the effector peptide, suggesting that the Q5 position is critical for substrate recognition and engagement by the catalytic domain. Previous experiments with a predicted Q5 interacting mutant in the paralog demethylase KDM5B support the importance of Q5-enzyme interaction for demethylation.^19^

**Figure 3:**
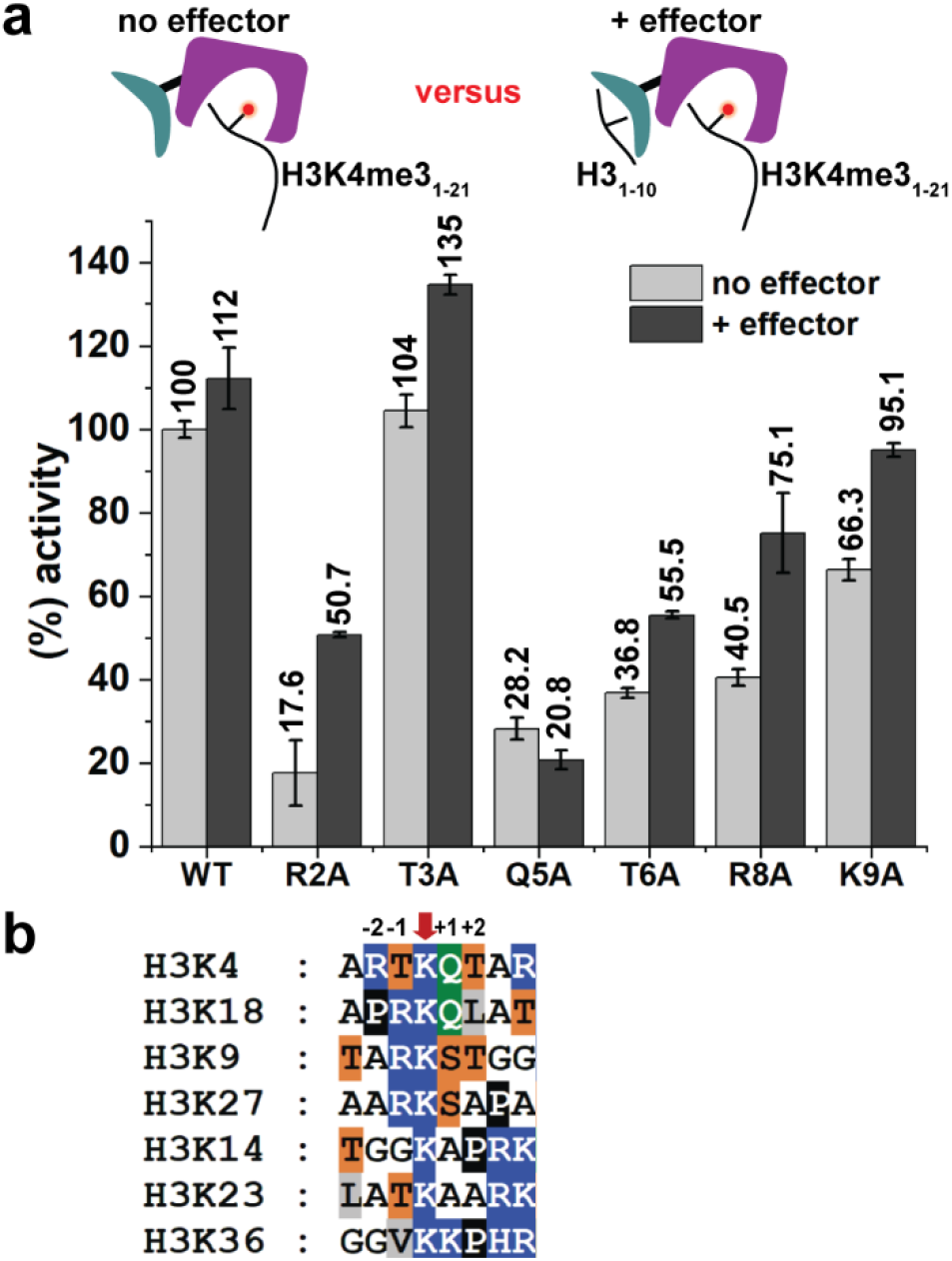
Effect of N-terminal H3 tail mutations on KDM5A activity. **a)** Impact of the PHD1 effector ligand on demethylase activity. KDM5A_1-797_ activity for 21mer H3K4me3 WT and mutant peptides in the absence and presence of 20 μM effector (unmodified 10mer peptide). Error bars represent the standard error of the mean of two independent experiments. **b)** Sequence alignment of H3 tail methylated lysine residues. Amino acids are color coded as follows: R/K/H in blue, T/S in orange, Q in green, A/G in white, P in black and L/V/I in grey.

In order to determine how these substrate mutants affect catalysis, we obtained apparent Michaelis-Menten kinetic constants under conditions where the PHD1 domain was saturated with effector peptide. While the measured k_cat_ was essentially the same for WT and mutant peptide substrates, the apparent K_M_ values were significantly different (Tables 1, S2 and Figures 3b, S1). Specifically, the K_M_^app^ for Q5A was over 30-fold higher than that for the WT peptide (415.2 ± 71.4 μM and 12.5 ± 0.9 μM, respectively), while the K_M_^app^ values for R2A, T6A and R8A were approximately 12-, 8- and 4-fold higher, respectively. We further probed the T6 interaction by mutating this residue to a serine and a valine. We found that the T6S peptide behaved like the WT substrate while substitution of T6 by valine resulted in a 3-fold increase in the K_M_. These results highlight the importance of the hydroxyl group of T6 in KDM5A-substrate interaction. Taken together, our findings suggest that R2, Q5, T6 and R8 all contribute to substrate recognition by KDM5A, with Q5 residue as the most important for substrate engagement. Interestingly, a comparison of our findings to those observed for the plant homolog JMJ14 points to species-specific substrate recognition in H3K4 demethylases. In the JMJ14 study, mutation of the residues that interact with H3R2 abrogated JMJ14 activity while H3T6 had no direct interaction with JMJ14.^19^ Such differences between KDM5s and the plant homolog are not surprising given differences in domain architecture as well as 50-60% similarity of their catalytic domains.

**Table 1.**
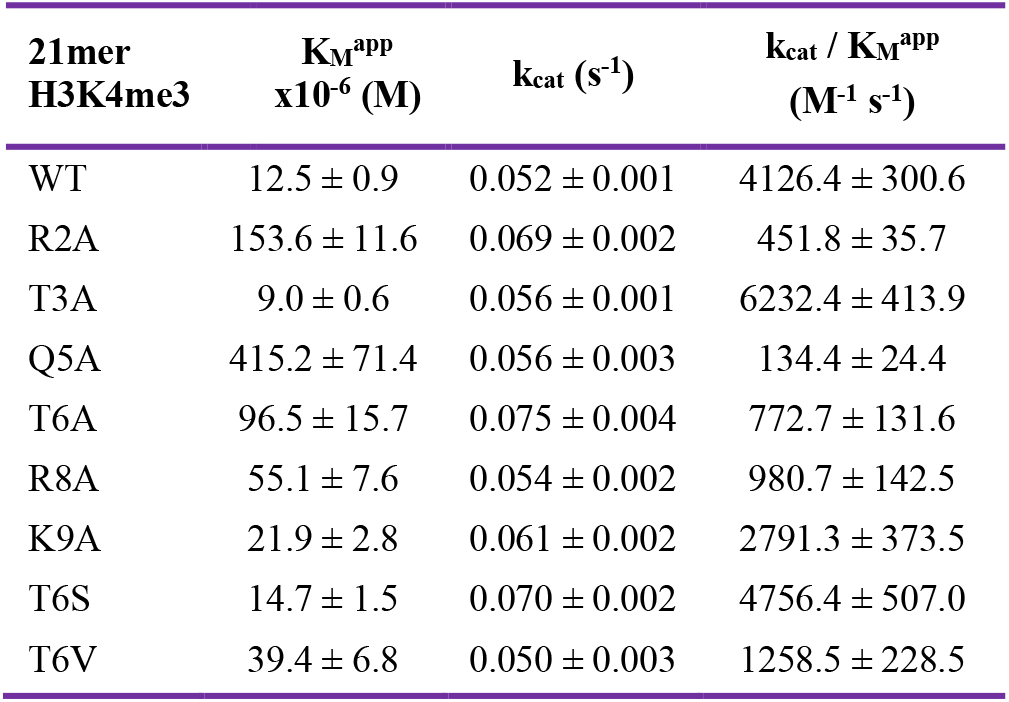
Apparent Michaelis-Menten kinetic parameters for 21mer H3K4me3 peptides in the presence of saturating (20μM) effector peptide (unmodified H3 10mer, aa 1-10). Results are means ± SEM of two independent experiments.

| 21mer H3K4me3 | $K_M^{app}$<br>$\times 10^{-6}$ (M) | $k_{cat}$ ($s^{-1}$) | $k_{cat} / K_M^{app}$<br>($M^{-1} s^{-1}$) |
| --- | --- | --- | --- |
| WT | 12.5 ± 0.9 | 0.052 ± 0.001 | 4126.4 ± 300.6 |
| R2A | 153.6 ± 11.6 | 0.069 ± 0.002 | 451.8 ± 35.7 |
| T3A | 9.0 ± 0.6 | 0.056 ± 0.001 | 6232.4 ± 413.9 |
| Q5A | 415.2 ± 71.4 | 0.056 ± 0.003 | 134.4 ± 24.4 |
| T6A | 96.5 ± 15.7 | 0.075 ± 0.004 | 772.7 ± 131.6 |
| R8A | 55.1 ± 7.6 | 0.054 ± 0.002 | 980.7 ± 142.5 |
| K9A | 21.9 ± 2.8 | 0.061 ± 0.002 | 2791.3 ± 373.5 |
| T6S | 14.7 ± 1.5 | 0.070 ± 0.002 | 4756.4 ± 507.0 |
| T6V | 39.4 ± 6.8 | 0.050 ± 0.003 | 1258.5 ± 228.5 |

A sequence alignment of the H3 tail lysine residues that undergo methylation (K9, K14, etc) (Figure 3b), together with our data, provides further insights into KDM5A specificity for H3K4. For the various methylated lysines, Arg in −2 (H3R2) position is replaced by a small aliphatic residue (alanine, glycine or proline), while Thr in +2 (H3T6) position is replaced by an aliphatic residue in all instances except K9. K9 however differs from K4 at the −2 (H3R2) and +1 (H3Q5) positions, substitutions sufficient to render H3K9me3 an unfavorable substrate for KDM5A.^7^ Finally, in addition to K4, the only H3 lysine that has a glutamine at position +1 (H3Q5 position) is K18. Since methylation of K18 is a newly detected mark with unknown function,^8, 10^ we tested whether KDM5A_1-797_ can demethylate K18. When H3K18me3 (aa 12-32) peptide was incubated with KDM5A_1-797_, no demethylation was detected (Figure S3). The observation that the adjacent glutamine is not sufficient for demethylation further supports recognition of an extended H3 region by KDM5A.

### KM5A recognizes an H3 epitope distal to K4

In addition to the N-terminal half of the 21mer H3K4me3 peptide substrate (aa A1-K9), we observed a significant reduction of KDM5A activity when P16 was substituted by alanine (~50%, Figure 2). As the PHD1 domain engages only the first 10 residues of H3 ^3–4^ and mutations in downstream positions are not expected to affect peptide engagement by PHD1, the decreased activity for P16A suggests that structural rigidity at position 16 may be important for substrate engagement. Furthermore, alanine substitution at the following position (R17A) decreased KDM5A activity to 62% (Figure 2). These findings led us to hypothesize that the basic residues surrounding P16 (K14, R17 and K18) may engage in interactions with KDM5A, and that their effect on substrate binding may be cumulative.

To test this hypothesis, we utilized peptide substrates that either lacked the K14-K18 basic patch (13mer peptide) or the three basic residues were mutated to alanines (K14A/R17A/K18A triple mutant, 18mer-AAA), and compared them to 18mer WT peptide substrate (Figure 4a). Both the deletion and alanine substitution of this basic patch substantially impaired demethylation of these substrates (approximately 28-35% compared to 18mer WT peptide). Further kinetic characterization showed that when this basic patch is removed, either by deletion or mutagenesis, the K_M_^app^ increases 8-fold compared to 18mer WT peptide (Tables 2, S3 and Figures 4b, S2). These results support our hypothesis that the K14-K18 basic patch is engaged by KDM5A and that P16 provides required structural rigidity for the basic patch-KDM5A interaction. The observed difference in K_M_^app^ also explains the preference of KDM5 enzymes towards longer peptide substrates.^3–4^

**Table 2.**
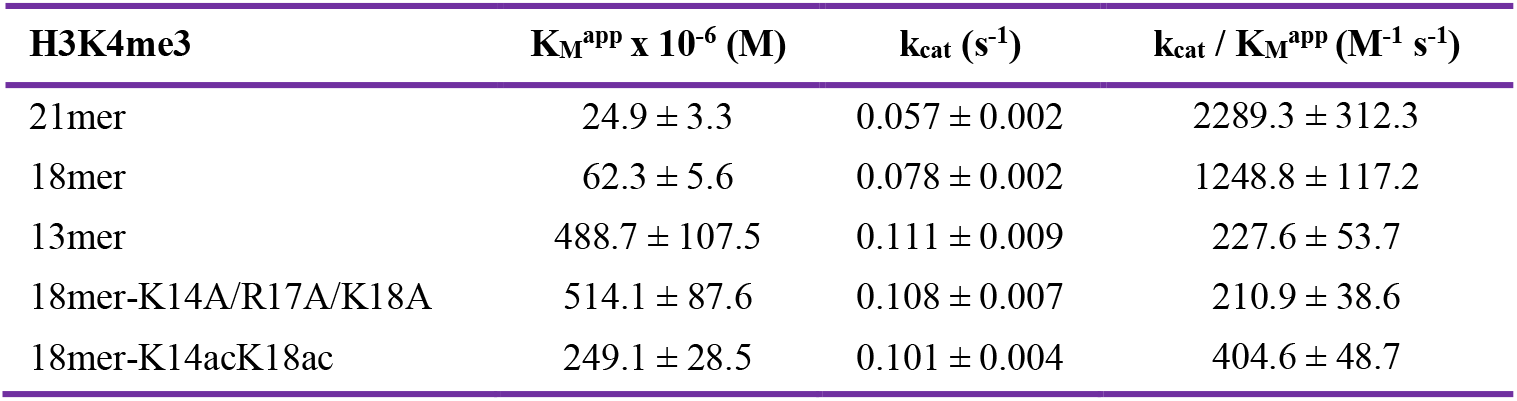
Apparent Michaelis-Menten kinetic parameters for H3K4me3 WT and mutant peptides of various lengths. Results are means ± SEM of two independent experiments.

**Figure 4:**
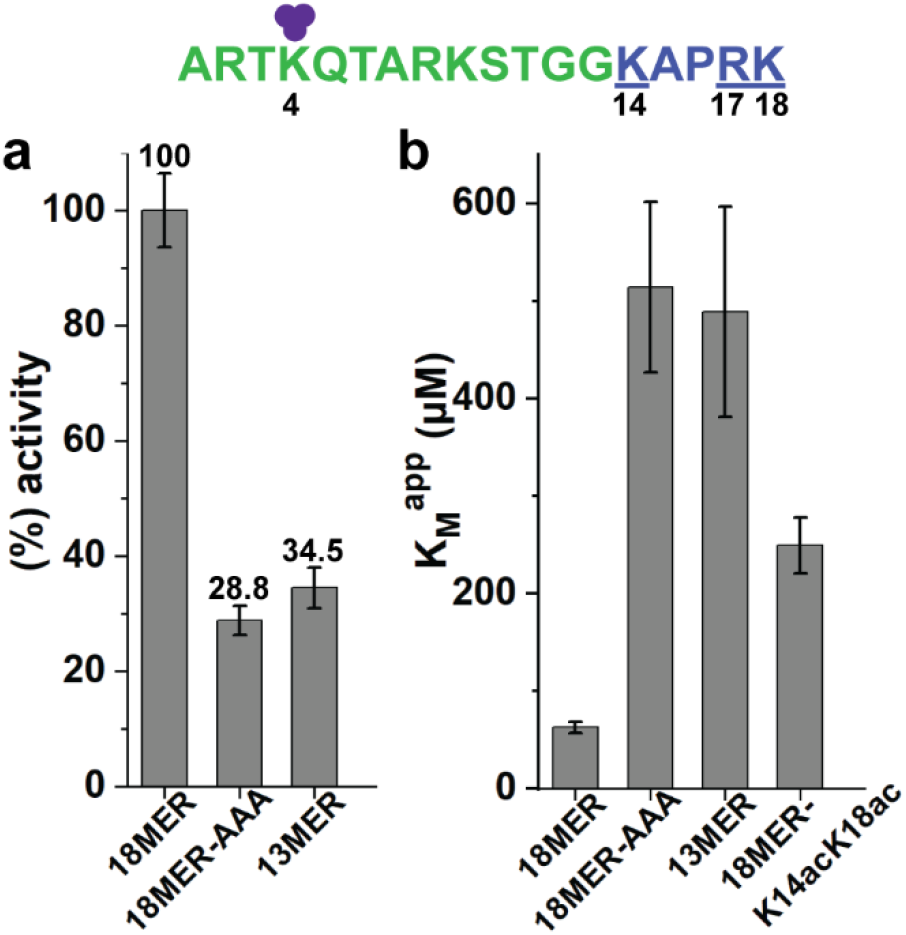
Recognition of H3 K14-K18 basic patch by KDM5A. **a)** KDM5A_1-797_ activity for H3K4me3 WT and basic patch mutant/deletion peptides normalized to the 18mer WT peptide reaction (considered as 100%). **b)** Apparent K_M_ values for the various basic patch peptides. Error bars represent the standard error of the mean of two independent experiments. (purple circles: methyl groups, green: 13mer, blue underlined: basic patch residues)

Acetylation of K14 and K18 is associated with active transcription, and a previous study identified a positive cross-talk between these modifications and K4me3.^8^ Hence, we proceeded to test how acetylation of K14 (K14ac) and K18 (K18ac) affects KDM5A substrate engagement and activity. Utilizing an 18mer peptide with acetylated lysines 14 and 18 (18mer-K14acK18ac), we found that acetylation of the basic patch increased the K_M_^app^ 4-fold relative to WT (Tables 2, S3 and Figures 4b, S2). This finding reveals that post-translational modifications installed on the K14-K18 basic patch can modulate KDM5A activity. Interestingly, K14ac was shown to inhibit K4me2 demethylation by histone demethylase KDM1A (LSD1).^20^

## CONCLUSION

Our investigations have defined substrate requirements for demethylation by human histone demethylase KDM5A. Specifically, we showed that engagement of H3Q5 is crucial for substrate binding, while H3-R2, T6 and R8 further contribute to H3K4me3-specific recognition by KDM5A. These findings provide a rationale for selectivity of KDM5A towards H3K4me3 mark over other methylated lysine residues in H3. Interestingly, we found that a basic patch on the histone tail downstream of H3K4 (aa K14-K18) is also involved in interactions with the enzyme. We demonstrated that post-translational modifications of this distal epitope can alter demethylase activity. To our knowledge, this is the first time that extended interactions between the catalytic domain of KDM5A and the H3 tail are probed, and a distal basic epitope on H3 is identified to modulate KDM5A demethylation. Combined with the allosteric regulation achieved through the PHD1 domain-effector peptide interaction, these findings provide important insights into how chromatin context can regulate catalytic activity of KDM5A, and consequently H3K4 methylation status.

## ASSOCIATED CONTENT

### Supporting Information

Experimental procedures, sequences of H3 peptides, M-M curves, K_M_ comparison graph and expanded M-M tables.

## AUTHOR INFORMATION

### Author Contributions

NP performed the experiments. JEL contributed to protein productions. The manuscript was written through contributions of all authors.

### Notes

The authors declare no competing financial interests.

## FOR TABLE OF CONTENTS ONLY

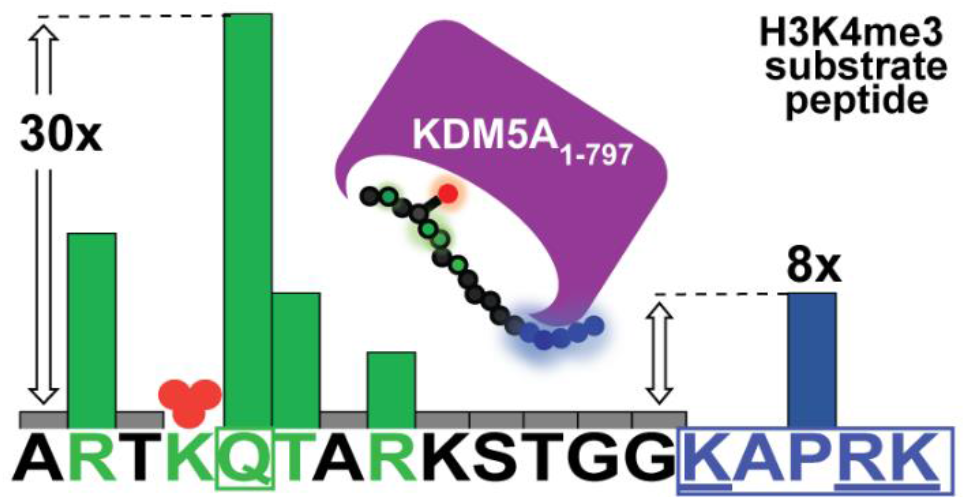

