## Supporting Information for "Extended Recognition of the Histone H3 Tail by Histone Demethylase KDM5A"

### Experimental Procedures

#### Expression and purification of recombinant $KDM5A_{1-797}$

The $KDM5A_{1-797}$ construct was expressed with an N-terminal polyhistidine tag in Sf21 cells following the Invitrogen Bac-to-Bac Baculovirus expression system protocol. The expression and purification of $KDM5A_{1-797}$ has been previously described.[1] Briefly, $KDM5A_{1–797}$ purified bacmid was transfected in Sf21 cells to obtain the P1 viral stock. To make P2, 20ml of $2x10^6$ cells/ml were infected with 2ml of P1 virus and incubated for 48–60 h. The cells were subsequently spun down and the supernatant was collected and sterile filtered to obtain P2 viral stock. The P2 viral stock was used to infect Sf21 cells for protein expression. Generally, 1L of Sf21 at $2x10^6$ cells per ml was infected with 40 ml of P2 virus for 48–56 h. Cells were then collected by centrifugation and resuspended in lysis buffer (25 mM HEPES pH 7.9, 350 mM NaCl, 5 mM KCl, 1.5 mM $MgCl_2$, 10 mM imidazole, 2 μg/ml aprotinin, 3 μg/ml leupeptin, 3 μg/ml pepstatin, 1 mM PMSF). Cells were lysed by multiple passages through an Emulsiflex cell homogenizer. Following centrifugation of the lysate, the supernatant was incubated with cobalt resin equilibrated in lysis buffer for 1 h at 4 °C. After incubation, the resin was washed with wash buffer (25 mM HEPES pH 7.9, 350 mM NaCl, 0.5 mM $MgCl_2$, 10% glycerol, 10 mM imidazole, 2 μg/ml aprotinin, 3 μg/ml leupeptin, 3 μg/ml pepstatin, 1 mM PMSF). His-$KDM5A_{1-797}$ was eluted with elution buffer (25 mM HEPES pH 7.9, 100 mM NaCl, 0.5 mM $MgCl_2$, 10% glycerol, 100 mM imidazole, 0.5 mM TCEP). Fractions containing protein of the highest purity (as determined by SDS-PAGE) were pooled and the polyhistidine affinity tag was removed by overnight incubation with TEV protease at 4 °C in the dialysis buffer (25 mM HEPES pH 7.9, 100 mM NaCl and 2 mM DTT). After cleavage, protein was further purified by size-exclusion chromatography (Superdex Hiload 200 26/60) in a buffer of 25 mM HEPES pH 7.5 and 50 mM KCl. Samples were concentrated using a 50,000-Da molecular weight cutoff Amicon centrifugal filter, flash-frozen in liquid nitrogen and stored at -80 °C.

#### KDM5A demethylation assay

Demethylation of methylated peptide substrates by $KDM5A_{1-797}$ was measured using the formaldehyde dehydrogenase (FDH) assay that reports on the release of by-product formaldehyde, which has been described previously.[2] All measurements were carried out at 21 °C in volumes of 100 μL dispensed in polystyrene nonbinding 96-well black flat-bottomed plates (Corning). The enzyme concentration was 1 μM in all experiments. The assay buffer used was 50 mM HEPES pH 7.5, 50 mM KCl, 50 μM ferrous ammonium sulfate ($(NH_4)_2Fe(SO_4)_2$), 1 mM alpha-ketoglutarate (α-KG), 2 mM ascorbate, 2 mM $NAD^+$ and 0.05 U FDH. $(NH_4)_2Fe(SO_4)_2$, ascorbate and α-KG were prepared fresh before each measurement, and $KDM5A_{1-797}$ was thawed in iced water. The reactions were initiated by addition of assay cocktail ($(NH_4)_2Fe(SO_4)_2$, ascorbate, $NAD^+$, FDH and $KDM5A_{1-797}$) into wells containing substrate cocktail (peptide and α-KG). The reactions were followed in 20-s intervals on a SpectraMax M5e (Molecular Devices) using 350 nm excitation and 460 nm emission wavelengths for a minimum of 10 min. Samples without peptide were included for each substrate and used for baseline correction and as negative controls. The amount of product formed per second was determined by using a NADH standard curve. Results are means ± SEM of two independent experiments (performed on different days with same enzyme preparation).

For the end-point assay, reactions of $KDM5A_{1-797}$ with 50 μM peptide were followed for 5 min. The activity of $KDM5A_{1-797}$ for the various peptide substrates is expressed as "percentage of activity" in respect to the wild-type H3K4me3 peptide reaction (considered as 100%).

For the Michaelis-Menten kinetics, various peptide concentrations were tested. Initial reaction velocities were calculated in Origin (OriginLab, Northampton, MA, U.S.A.) by obtaining the slope from the best-fit line for a time course of 2.3 min. Based on the initial reaction rates, the apparent $K_M$ and $V_{max}$ values were determined using the Michaelis−Menten function of Origin. Because the peptide substrates can also bind to the PHD1 domain and enhance substrate binding to the catalytic domain of KDM5A,[1, 3] true Michaelis-Menten kinetics will not be observed, and values of $K_M^{app}$ are consequently reported.

When the assays were performed in the presence of effector peptide (unmodified H3 10mer), 20 μM ($20xK_D$)[3] effector was included in the assay cocktail.

**Supplementary Table S1:** Sequences of histone H3 peptides used in this study.

| Peptide | Sequence |
|---|---|
| **21mer-H3K4me3-WT** | ARTK(me3)QTARKSTGGKAPRKQLA |
| **21mer-H3K4me3-R2A** | A**A**TK(me3)QTARKSTGGKAPRKQLA |
| **21mer-H3K4me3-T3A** | AR**A**K(me3)QTARKSTGGKAPRKQLA |
| **21mer-H3K4me3-Q5A** | ARTK(me3)**A**TARKSTGGKAPRKQLA |
| **21mer-H3K4me3-T6A** | ARTK(me3)Q**A**ARKSTGGKAPRKQLA |
| **21mer-H3K4me3-T6S** | ARTK(me3)Q**S**ARKSTGGKAPRKQLA |
| **21mer-H3K4me3-T6V** | ARTK(me3)Q**V**ARKSTGGKAPRKQLA |
| **21mer-H3K4me3-R8A** | ARTK(me3)QTA**A**KSTGGKAPRKQLA |
| **21mer-H3K4me3-K9A** | ARTK(me3)QTAR**A**STGGKAPRKQLA |
| **21mer-H3K4me3-S10A** | ARTK(me3)QTARK**A**TGGKAPRKQLA |
| **21mer-H3K4me3-T11A** | ARTK(me3)QTARKS**A**GGKAPRKQLA |
| **21mer-H3K4me3-G12A** | ARTK(me3)QTARKST**A**GKAPRKQLA |
| **21mer-H3K4me3-G13A** | ARTK(me3)QTARKSTG**A**KAPRKQLA |
| **21mer-H3K4me3-K14A** | ARTK(me3)QTARKSTGG**A**APRKQLA |
| **21mer-H3K4me3-P16A** | ARTK(me3)QTARKSTGGKA**A**RKQLA |
| **21mer-H3K4me3-R17A** | ARTK(me3)QTARKSTGGKAP**A**KQLA |
| **21mer-H3K4me3-K18A** | ARTK(me3)QTARKSTGGKAPR**A**QLA |
| **21mer-H3K4me3-Q19A** | ARTK(me3)QTARKSTGGKAPRK**A**LA |
| **21mer-H3K4me3-L20A** | ARTK(me3)QTARKSTGGKAPRKQ**A**A |
| **18mer-H3K4me3-WT** | ARTK(me3)QTARKSTGGKAPRK |
| **13mer- H3K4me3-WT** | ARTK(me3)QTARKSTGG |
| **18mer-H3K4me3-AAA (K14A/R17A/K18A)** | ARTK(me3)QTARKSTGG**A**AP**AA** |
| **18mer-H3K4me3-K14acK18ac** | ARTK(me3)QTARKSTGG**K(ac)**APR**K(ac)** |
| **21mer-H3K18me3 (aa 12-32)** | GGKAPRK(me3)QLATKAARKSAPAT |
| **Effector (10mer-H3K4me0)** | ARTKQTARKS |

**Supplementary Table S2.** Expanded table for apparent Michaelis-Menten kinetic parameters for 21mer H3K4me3 peptides in the presence of saturating (20μM) effector peptide (unmodified H3 10mer, aa 1-10). Results are means ± SEM of two independent experiments.

| 21mer H3K4me3 | $K_M^{app}$ x $10^{-6}$ (M) | $k_{cat}$ ($s^{-1}$) | $k_{cat}$ / $K_M^{app}$ ($M^{-1}$ $s^{-1}$) | Fold change* in $K_M^{app}$ | Fold change* in $k_{cat}$ / $K_M^{app}$ |
|---|---|---|---|---|---|
| WT | 12.5 ± 0.9 | 0.052 ± 0.001 | 4126.4 ± 300.6 | 1.0 ± 0.1 | 1.0 ± 0.1 |
| R2A | 153.6 ± 11.6 | 0.069 ± 0.002 | 451.8 ± 35.7 | 12.3 ± 1.3 | 0.11 ± 0.01 |
| T3A | 9.0 ± 0.6 | 0.056 ± 0.001 | 6232.4 ± 413.9 | 0.72 ± 0.07 | 1.5 ± 0.2 |
| Q5A | 415.2 ± 71.4 | 0.056 ± 0.003 | 134.4 ± 24.4 | 33.2 ± 6.2 | 0.033 ± 0.006 |
| T6A | 96.5 ± 15.7 | 0.075 ± 0.004 | 772.7 ± 131.6 | 7.7 ± 1.4 | 0.19 ± 0.03 |
| R8A | 55.1 ± 7.6 | 0.054 ± 0.002 | 980.7 ± 142.5 | 4.4 ± 0.7 | 0.24 ± 0.04 |
| K9A | 21.9 ± 2.8 | 0.061 ± 0.002 | 2791.3 ± 373.5 | 1.8 ± 0.3 | 0.7 ± 0.1 |
| T6S | 14.7 ± 1.5 | 0.070 ± 0.002 | 4756.4 ± 507.0 | 1.2 ± 0.1 | 1.2 ± 0.1 |
| T6V | 39.4 ± 6.8 | 0.050 ± 0.003 | 1258.5 ± 228.5 | 3.2 ± 0.6 | 0.31 ± 0.06 |

* The fold change was calculated by dividing the mutant value by the WT value.

**Supplementary Table S3.** Expanded table for apparent Michaelis-Menten kinetic parameters for H3K4me3 WT and mutant peptides of various lengths. Results are means ± SEM of two independent experiments.

| **Peptide H3K4me3** | **$K_M^{app}$ x $10^{-6}$ (M)** | **$k_{cat}$ ($s^{-1}$)** | **$k_{cat}$ / $K_M^{app}$ ($M^{-1}$ $s^{-1}$)** | **Fold change* in $K_M^{app}$** | **Fold change* in $k_{cat}$ / $K_M^{app}$** |
|---|---|---|---|---|---|
| 21mer | 24.9 ± 3.3 | 0.057 ± 0.002 | 2289.3 ± 312.3 | 0.40 ± 0.06 | 1.8 ± 0.3 |
| 18mer | 62.3 ± 5.6 | 0.078 ± 0.002 | 1248.8 ± 117.2 | 1.0 ± 0.1 | 1.0 ± 0.1 |
| 13mer | 488.7 ± 107.5 | 0.111 ± 0.009 | 227.6 ± 53.7 | 7.8 ± 1.9 | 0.18 ± 0.05 |
| 18mer-K14A/R17A/ K18A | 514.1 ± 87.6 | 0.108 ± 0.007 | 210.9 ± 38.6 | 8.3 ± 1.6 | 0.17 ± 0.03 |
| 18mer-K14acK18ac | 249.1 ± 28.5 | 0.101 ± 0.004 | 404.6 ± 48.7 | 4.0 ± 0.6 | 0.32 ± 0.05 |

* The fold change was calculated by dividing the mutant value by the 18mer WT value.

**Supplementary Figure S1:** Michaelis-Menten kinetic curves for 21mer H3K4me3 WT and mutant peptides in the presence of 20µM (20x$K_D$) effector (unmodified 10mer peptide). Results are means ± SEM of two independent experiments.

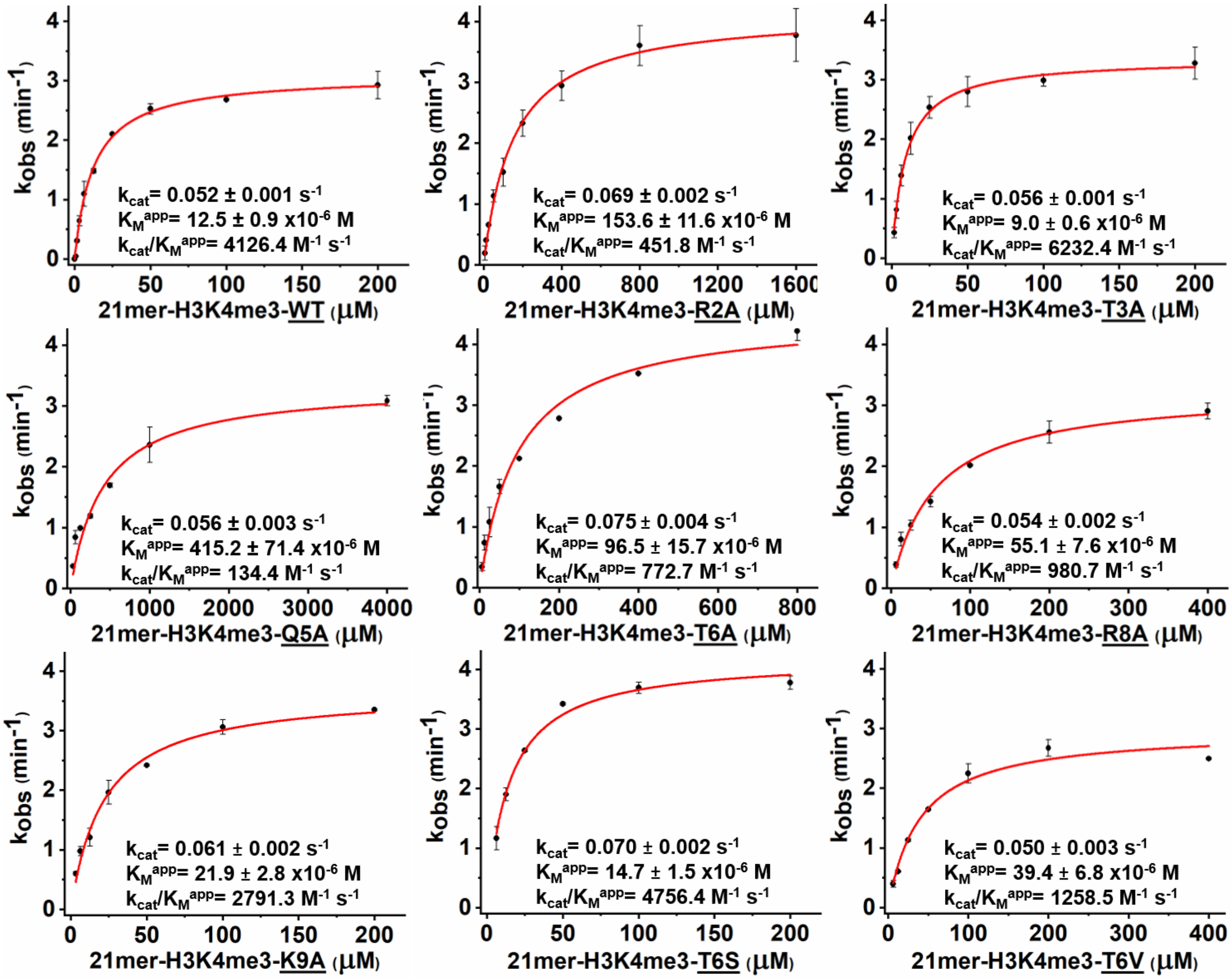

**Supplementary Figure S2:** Michaelis-Menten kinetic curves for H3K4me3 WT and mutant peptides of various lengths. The 21mer peptide sequence is also shown (purple circles: methyl groups, green: 13mer, blue underlined: “basic patch” residues that were mutated to Ala). Results are means ± SEM of two independent experiments.

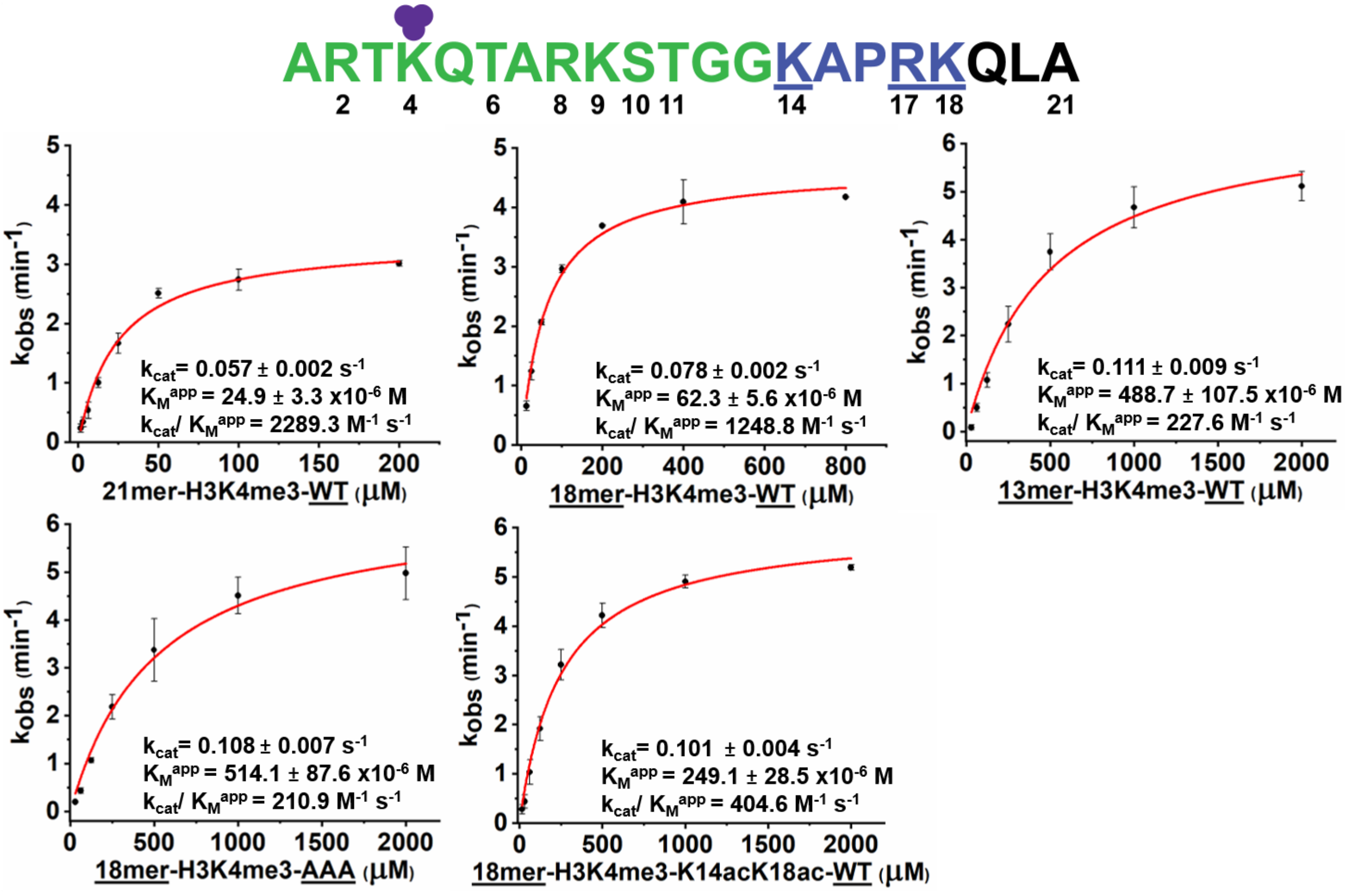


**Supplementary Figure S3: KDM5A1-797 does not demethylate H3K18me3.** Formaldehyde dehydrogenase assay reading for 21mer H3K4me3 and H3K18me3 peptides. No change in fluorescence was observed when H3K18me3 (aa 12-32) was provided as substrate.

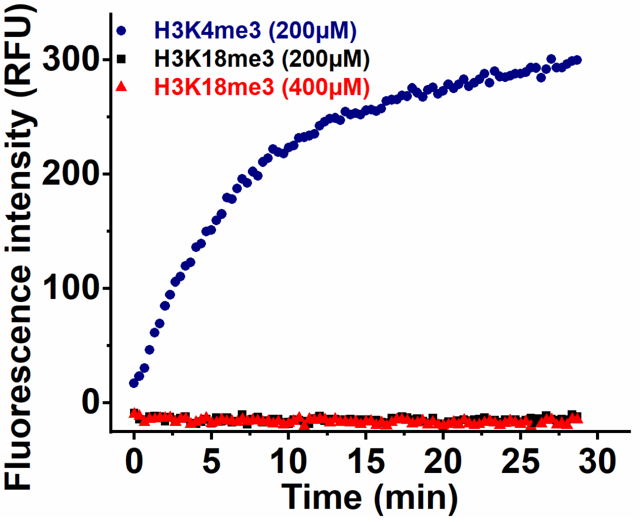

**Figure S4.** Apparent $K_M$ values for 21mer H3K4me3 WT and mutant peptides in the presence of 20 μM effector. Error bars represent the standard error of the mean of two independent experiments.

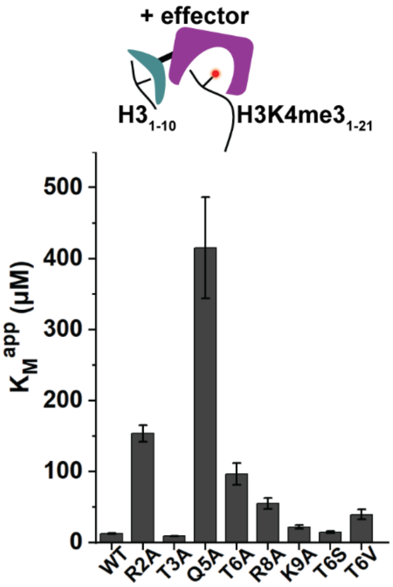
